# Cytoskeletal engineering through Formin-like 1 overexpression enhances T cell infiltration and antitumor potency in solid tumors

**DOI:** 10.64898/2026.08.20.744715

**Authors:** Jeffrey W. Chung, Jessica Olivas-Corral, Ashley M. Wood, Heidi Solis, Ashton L. Sigler, Edward Ning, Michelle E. Allen, Kayla H. Thompson, Jordan Jacobelli

## Abstract

Solid tumors are often surrounded by abnormal vasculature and a dense collagen-rich extracellular matrix that severely restrict the infiltration of T cells, including tumor-infiltrating lymphocytes (TILs) and chimeric antigen receptor (CAR)-T cells. These physical barriers represent a major obstacle to the efficacy of adoptive T cell therapies in solid tumors. We previously identified Formin-like 1 (FMNL1) as a cytoskeletal regulator critical for T cell extravasation and migration through restrictive environments, making it a promising target to improve T cell infiltration into tumors. Here, we developed a bioengineering platform to enhance T cell cytoskeletal dynamics by overexpressing FMNL1 in TILs and CAR-T cells. FMNL1 overexpression significantly increased T cell migration through restrictive pores in transwell assays, supporting enhanced migratory capacity of T cells under mechanically constraining conditions. Importantly, FMNL1 overexpression did not impair T cell reactivation or cytotoxic function *in vitro*. In murine models of melanoma and lung carcinoma characterized by limited effector T cell infiltration, FMNL1-overexpressing TILs and CAR-T cells had significantly increased accumulation at tumor sites compared to controls. Importantly, enhanced tumor accumulation resulted in improved therapeutic activity, as adoptive transfer of FMNL1-overexpressing CAR-T cells limited tumor growth and prolonged the survival of tumor-bearing mice in multiple melanoma models. Together, our findings identify FMNL1 as a broadly applicable cytoskeletal engineering target to enhance T cell accumulation and persistence in restrictive tumor microenvironments, thereby overcoming a fundamental limitation of adoptive cellular immunotherapy in solid tumors.

## Introduction

One of the major challenges for targeting solid tumors with adoptive cell therapies, including tumor infiltrating lymphocytes (TILs) and chimeric antigen receptor (CAR)-T cells, is the inability of T cells to sufficiently migrate and accumulate in the tumor tissue [1, 2]. Several mechanisms are used by solid tumors to prevent the migration of cytotoxic T cells into the tumor parenchyma. For instance, the aberrant vasculature around the tumor can impede T cell homing to solid tumors [3]. For tumors to rapidly grow, they induce neo-angiogenesis with cytokines like vascular endothelial growth factor (VEGF) and chemokines such as VEGF-correlated chemokine-1 [4, 5]. As angiogenesis becomes dysregulated, the tumor vasculature becomes leaky and irregular in size and structure, creating an environment that is hypoxic, nutrient-depleted, and has irregular blood flow reducing efficient T cell extravasation [3, 5, 6].

Additionally, the increase in angiogenesis can cause endothelial cells to downregulate the expression of adhesion molecules such as ICAM-1 and VCAM-1 which are required for T cell adhesion and extravasation [6]. Furthermore, the abnormal tumor vasculature may not express sufficient levels of chemokines to traffic T cells to the tumor [7], and the tumor microenvironment (TME) can also release chemokines that signal T cells to migrate away from the tumor [8].

Another major barrier for T cell migration into solid tumors is the extracellular matrix (ECM) surrounding the tumor. The ECM goes through dynamic chemical and biophysical changes during tumorigenesis characterized by dense deposits of fibrillar collagen and other components such as fibronectin. The release of TGFβ from stromal cells induces fibroblast activation and excessive production of ECM molecules, reducing ECM pore size and increasing the stiffness of the ECM. These environmental changes create a physical barrier that inhibits T cells from efficiently penetrating the tumor mass and targeting malignant cells [9–14].

A large body of research is exploring different strategies to enhance T cell effectiveness in solid tumors. Promising immune-based therapies include immune checkpoint inhibitors that block CTLA-4 and/or the PD-1/PD-L1 axis. Blocking these negative regulators of T cell activation can improve antitumor TIL and CAR-T function [15–17]. However, checkpoint inhibitor treatment fails in most patients that have poor T cell infiltration into solid tumors [2, 16, 18, 19]. Other studies have shown that regulation of angiogenesis through inhibiting VEGF signaling with antibodies normalizes blood vessels and induces adhesion molecules on endothelial cells [6, 20]. Others have engineered the expression of chemokine receptors or modulated atypical chemokine receptors to enhance T cell migration to tumors [21–26]. With the heterogenicity of solid tumors, these approaches unfortunately often have limited applicability as vascular environments and chemokine expression differ between tumor types [27, 28]. Additionally, these approaches do not address the challenge of anti-tumor T cell migration through dense ECM.

The actin cytoskeleton provides the driving force for T cells to migrate and interact with their environment. Formin-like-1 (FMNL1) and mammalian Diaphanous-related protein 1 (mDia1) are the two highly expressed formins in T cells [29–31]. FMNL1 and mDia1 are key cytoskeletal effector proteins that influence cytoskeletal dynamics by modulating actin polymerization, actin filament assembly, and protrusion formation. FMNL1 has an important role in T cell diapedesis and effector T cell trafficking to inflamed tissues by promoting transmigration of the nucleus through restrictive endothelial cell barriers [32]. FMNL1 physically associates with the nucleus and assists in the deformation of the T cell nucleus during migration through confined environments [29]. mDia1 promotes motility within 3D environments in conjunction with Myosin-II activity and can also regulate T cell trafficking [29, 33, 34]. Furthermore, FMNL1 has been found to be strongly expressed in immune cells infiltrating human hepatocellular carcinoma [35]. This elevated FMNL1 expression in immune cells was associated with immune-inflamed (“hot”) tumor microenvironments, enhanced responses to immune checkpoint inhibitor therapy, and improved patient prognosis [35], suggesting a potential role for FMNL1 in promoting effective antitumor immunity. Given these findings and the importance of FMNL1 and mDia1 in the ability of T cells to home to inflamed sites and migrate through restrictive interstitial tissues, we hypothesized that overexpression of FMNL1 or mDia1 would enhance the ability of TILs and CAR-T cells to migrate and accumulate at solid tumor sites *in vivo*.

## Methods

### Mice

All experiments were performed in compliance with the study protocol approved by the University of Colorado Anschutz Medical Campus Institutional Animal Care and Use Committee (IACUC, protocol no: 00937). The study was conducted in accordance with the local legislation and institutional requirements. CD45.1 congenically marked C57BL/6 mice (purchased from Charles River, strain #564) and Rag knock-out mice (purchased from The Jackson Laboratory, strain #002216) were used as tumor recipients. Wild-type (WT), OT-I, and pmel-1 TCR transgenic mice were bred on the C57BL/6 background. OT-I mice are a TCR transgenic strain specific to SIINFEKL peptide of ovalbumin (Ova 257-264). Pmel-1 TCR transgenic mice are specific for premelanosome protein pmel-17/gp100. TIL experiments used OT-I or pmel-1 mice as T cell donors. CAR-T experiments used CD8 T cells isolated from WT C57BL/6 mice. All mice used in experiments were 6-16 weeks old. T cell donor mice were sex-matched with the tumor-bearing recipients. All efforts were made to minimize mouse suffering.

### Flow cytometry

T cells were analyzed and/or sorted via flow cytometry on an Aurora (Cytek Biosciences) spectral flow cytometer or a Bio-Rad S3e cell sorter. All flow cytometry data were analyzed with FlowJo (FloJo) software and/or SpectroFlo (Cytek Biosciences). The following antibodies/dyes were used throughout this study:

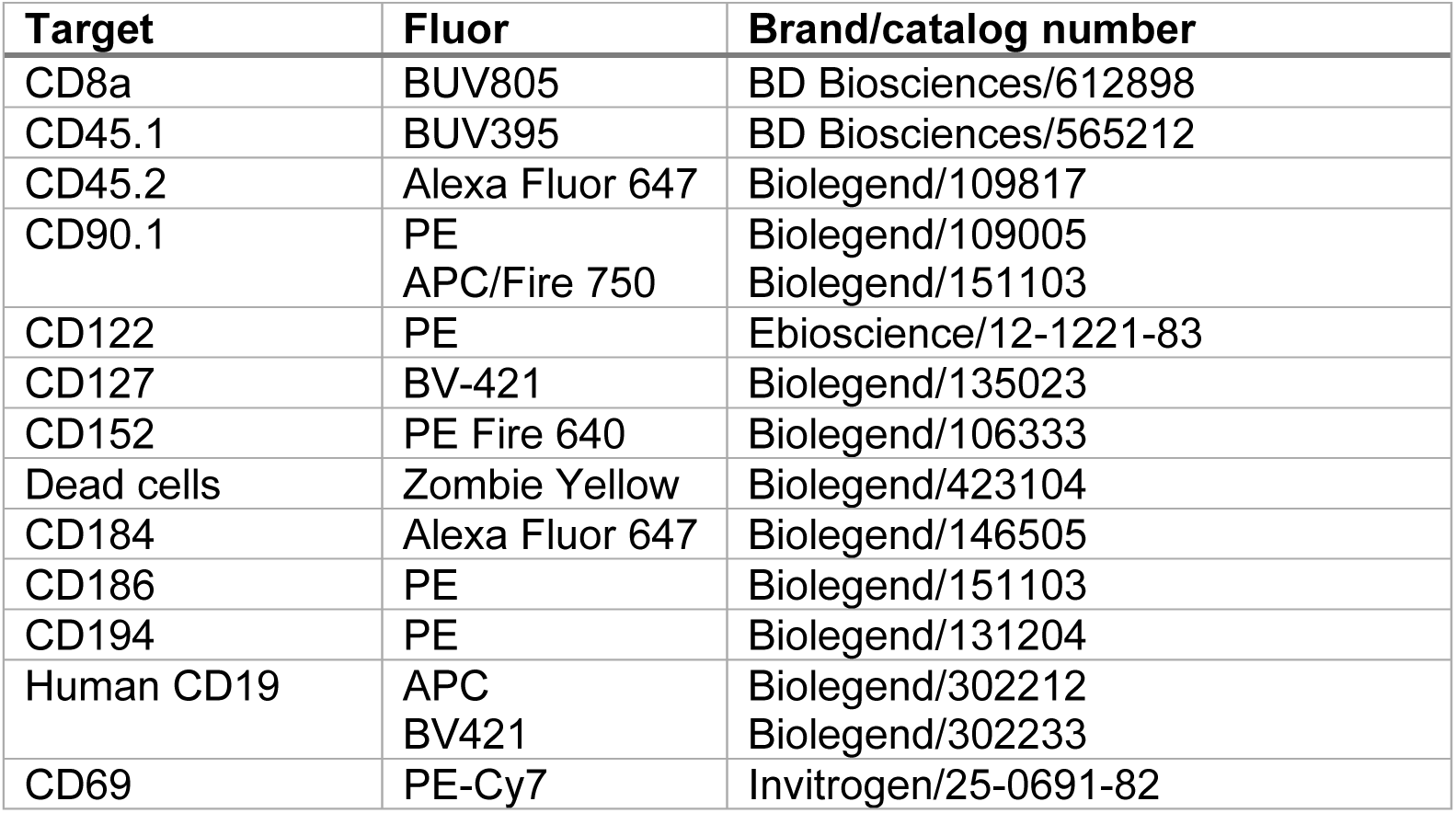

| Target | Fluor | Brand/catalog number |
| --- | --- | --- |
| CD8a | BUV805 | BD Biosciences/612898 |
| CD45.1 | BUV395 | BD Biosciences/565212 |
| CD45.2 | Alexa Fluor 647 | Biolegend/109817 |
| CD90.1 | PE<br>APC/Fire 750 | Biolegend/109005<br>Biolegend/151103 |
| CD122 | PE | Ebioscience/12-1221-83 |
| CD127 | BV-421 | Biolegend/135023 |
| CD152 | PE Fire 640 | Biolegend/106333 |
| Dead cells | Zombie Yellow | Biolegend/423104 |
| CD184 | Alexa Fluor 647 | Biolegend/146505 |
| CD186 | PE | Biolegend/151103 |
| CD194 | PE | Biolegend/131204 |
| Human CD19 | APC<br>BV421 | Biolegend/302212<br>Biolegend/302233 |
| CD69 | PE-Cy7 | Invitrogen/25-0691-82 |

### Cell culture

OT-I and pmel-1 T cells were cultured in R10 media: RPMI 1640 (Corning) supplemented with L glutamine (292 µg/mL), penicillin (100 U/mL), streptomycin (100 µg/mL), and β-mercaptoethanol (50µM) (all purchased from Gibco) and 10% Fetal Bovine Serum (FBS). The Plat-E cell line (Cell Biolabs) was cultured in D10 media: DMEM (Corning) supplemented with 10mM HEPES (Corning), L-glutamine, penicillin, streptomycin, and β-mercaptoethanol and 10% FBS. CAR-T cells were cultured in R10 media enriched with the following additions: sodium pyruvate (100 mM), MEM non-essential amino acids (1%), and GlutaMAX (all purchased from Gibco). Lewis lung carcinoma cell line LL/2 (CRL-1642) was obtained from ATCC. The B16 and B78ChOva melanoma cells were a gift from Dr. Matthew Krummel (UCSF). The B78ChOva melanoma cell line was generated via transfection of B78 melanoma cells derived from a B16 sub-line that lost melanin expression with a vector containing mCherry and chicken ovalbumin (Ova) [36, 37]. All cell lines were cultured in DMEM with 10% heat-inactivated FBS in a humidified atmosphere of 10% CO_2_.

### T cell isolation, activation, and culture

Lymphocytes were isolated from dissociated lymph nodes (LN) and spleen from donor mice. Spleens were mashed through 100 µm strainers into 175 mM ammonium chloride to lyse red blood cells and then spun down and washed with R10. For TIL experiments using OT-I T cells, T cells were stimulated with splenocytes pulsed with 100 ng/ml SIINFEKL peptide for 30 minutes and washed 3 times. OT-I T cells were plated in 24 well plates with peptide-pulsed splenocytes at a 1:2 ratio at 10^6^ cells/mL in R10 media. Alternatively, OT-I T cells, or WT T cells, were activated with plate bound CD3 and CD28 at 2×10^6^ cells/mL. Cells were fed at 48 hours with fresh media and 10 U/mL IL-2 (AIDS Research and Reference Reagent Program, Division of AIDS, National Institute of Allergy and Infectious Diseases, National Institutes of Health, from M. Gately, Hoffmann-La Roche).

For experiments using pmel-1 T cells and CAR-T cells, total CD8 T cells were isolated using magnetic negative selection kits (Stemcell Technologies) according to manufacturer instructions. Isolated T cells were activated with Dynabeads Mouse T-Activator CD3/CD28 (thermoFisher; 11452D) at a 1:1 T cell to bead ratio. T cells were plated in a 24 well plate at 1 million/ml/well supplemented with 40 U/mL IL-2 and 10 µg/ml IL-7 (R&D, 207-IL-005). Dynabeads were removed 72 hours later.

### FMNL1 or mDia1 overexpression and CAR transduction

For viral vector supernatant production, 3×10^6^ Plat-E cells (a retroviral packaging cell line) were plated in 10 cm dishes and incubated at 37°C with CO_2_. The following day, the media was swapped with fresh media with 50 µM Chloroquine (Sigma). A calcium phosphate-DNA particle mixture was prepared as follows: 20 µg of retroviral vector, 62.5 µL of 2M CaCl_2_, and 5 µg pCL-Eco plasmid were mixed in water for a final volume of 500 µL and then combined with 500 µL 2x HBS. The 1000 µL calcium phosphate-DNA mixture was added to the Plat-E dish dropwise and incubated overnight. The Chloroquine media was then removed and replaced with fresh media. After another 24 hours, the supernatant containing the retrovirus was collected and used for T cell transductions. Fresh media was added to the Plat-E dish and the viral supernatant was collected 24 hours later for a second round of T cell transductions.

For the transduction, T cells were activated (as described above) and then 24-48 hours post-activation, they were transduced with Maloney murine leukemia retrovirus (MMLV) vectors expressing either FMNL1 and GFP under an internal ribosome entry site (IRES) or fluorescent protein only (e.g. Cerulean or TurboRFP). Cells were transduced a second time with the same viral vectors 24 hours later. On day 4 post activation, fluorescent protein marker-positive T cells were sorted using Bio-Rad S3e or Cytek Biosciences Aurora cell sorters. After 24 hours post sort, cells were transferred into recipient mice or used for *in vitro* assays. For CAR-T cell experiments, a murine stem cell virus (MSCV) retroviral vector expressing a CAR construct against human CD19 (a gift from Dr. James Scott-Browne, National Jewish Health) was transduced in combination with the MMLV FMNL1, MMLV mDia1, or the MMLV control constructs. T cells were analyzed for CAR+ by expression of CD90.1 included in the CAR vector [38].

### Generation of human CD19 expressing tumor cell lines

The tumor cell lines LL/2 and B78ChOva were stably transduced with lentivirus encoding human CD19 antigen (hCD19) (pLV.hEF1a.cHS4 hCD19t.WPRE a gift from Dr. Terry Fry, University of Colorado). Tumor cells were sorted for hCD19 expression using the Bio-Rad S3e cell sorter and validated for stable hCD19 expression.

### *In vitro* T cell restimulation and tumor killing assay

Control or FMNL1 overexpressing OT-I T cells were co-cultured with B78ChOva (mCherry+) tumor cells using a 6:1 effector:target cell ratio in a 96 well plate. Twenty-four hours later, T cells were removed and stained with CD69 to assess for activation by flow cytometry.

Percent tumor cells killed was calculated by quantifying tumor cell numbers after 24 hours of T cell-tumor cell co-culture and comparing to a no T cell tumor cell culture.

### Western blotting

FMNL1 was detected using a rabbit polyclonal antibody (Invitrogen; PA5-52516). As a loading control, α-tubulin (Millipore-Sigma; B-1-5-2) or GAPDH (Santa Cruz; sc-365564) was detected using a monoclonal mouse antibody. Antibody staining was visualized using the Azure Biosystems Sapphire Imager with secondary antibodies against mouse (Licor; 926-68072) and rabbit (Licor; 926-32213) antibodies. Band intensities were quantified by densitometry using the Sapphire Biomolecular Imager and associated software (Azure Biosystems).

### Ectopic tumor inoculation

Tumor cell lines listed above were cultured in DMEM medium with 10% FBS and passaged several times before inoculation. On the day of injection, tumor cells were trypsinized and washed with 10 mL of phosphate buffered saline (PBS) 3 times. Cells were resuspended at the indicated concentrations with/without the addition of Matrigel (Corning; 356231) as follows:

| Cell line | Cell concentration per mL | With Matrigel? |
| --- | --- | --- |
| LL/2-hCD19 | $10 \times 10^6$ | No |
| B16-F10 | $1.6 \times 10^6$ | Yes |
| B16-Ova | $1.6 \times 10^6$ | Yes |
| B78ChOva | $2 \times 10^6$ | Yes |
| B78ChOva-hCD19 | $3 \times 10^6$ | Yes |

Tumor cells were loaded into 28G needle syringes and 50 µL was injected subcutaneously into the flanks of CD45.1 C57BL/6 mice or Rag knock-out C57BL/6 mice. Tumor measurements were recorded with a manual caliper and tumor area was calculated in millimeters squared (length x width).

### Transwell assay

Wells of a 24-well plate were prepared with 3 µm transwell inserts (Corning; 3415) and RPMI supplemented with 0.5% FBS with 1 µg/mL CXCL12 (Peprotech) in the bottom chamber. 1 x 10^5^ control and FMNL1 overexpressing T cells were added to the top chambers and allowed to migrate for 1 hr at 37°C into the lower well. As a control, 2 x 10^4^ cells (20% of input cells added to transwells) were placed directly into bottom wells with no transwell as a standard to calculate the percentage of migrated cells. Migrated T cells were collected from the bottom wells and quantified for a fixed volume using a flow cytometer (Cytek Aurora).

### Tissue processing and analysis

For T cell trafficking experiments, on day −5 tumor cells were inoculated into recipient mice. On the same day T cells were collected from donor mice and were activated *ex vivo*, transduced with a control plasmid or FMNL1/ mDia1 expressing plasmid (as described above), and T cells were sorted on day −1. On day 0, control T cells and formin overexpressing T cells were mixed at a 1:1 ratio (1-5 x 10^5^ cells of each) and intravenously injected into tumor bearing mice. Two to six days post T cell transfer, mice were euthanized by CO_2_ inhalation. Blood was collected by cardiocentesis, and red blood cells were lysed for 60 minutes on ice with 175mM ammonium chloride. Spleen was collected and mechanically dissociated, and red blood cells were further lysed for 5 min at room temperature in 175mM ammonium chloride. Inguinal lymph nodes were collected and mechanically dissociated. As nondraining lymph nodes, the axillary and brachial lymph nodes were collected and mechanically dissociated. Tumors were carefully excised and mechanically dissociated. The dissociated tumors were then digested in collagenase D (0.786 Wunsch U/mL, Roche) and DNaseI (250 µg/mL, Roche) for 30 minutes at 37°C while rotating. Tumor cells were then filtered through a 100µM filter. After preparations of single cell suspensions, cells were stained with fluorescently labeled antibodies and analyzed by flow cytometry. Transferred T cells were identified as Live, CD8+, CD45.2+, GFP+, TurboRFP+, or Cerulean+. For CAR-T experiments, CD90.1+ was also used to identify transferred CAR-T cells. Only tumors containing ≥10 transferred T cells from either the control or FMNL1-overexpressing group were included in the analysis. Tumors with <10 cells of both populations were excluded to reduce variability from extremely low cell counts and ensure robust quantification.

### Multi-spectral analysis of tumor sections

Similar to the T cell tumor accumulation experiments, control and FMNL1-overexpressing OT-I T cells were co-transferred into B16-Ova tumor bearing mice, then 3-6 days post-T cell transfer the tumors were carefully excised, formalin-fixed and paraffin-embedded. In collaboration with the Human Immune Monitoring Shared Resource (HIMSR) at the University of Colorado School of Medicine we performed multi-spectral imaging of tumor sections using the Lunaphore COMET imaging platform (Bio-techne). Tissue sections were stained for the following markers:

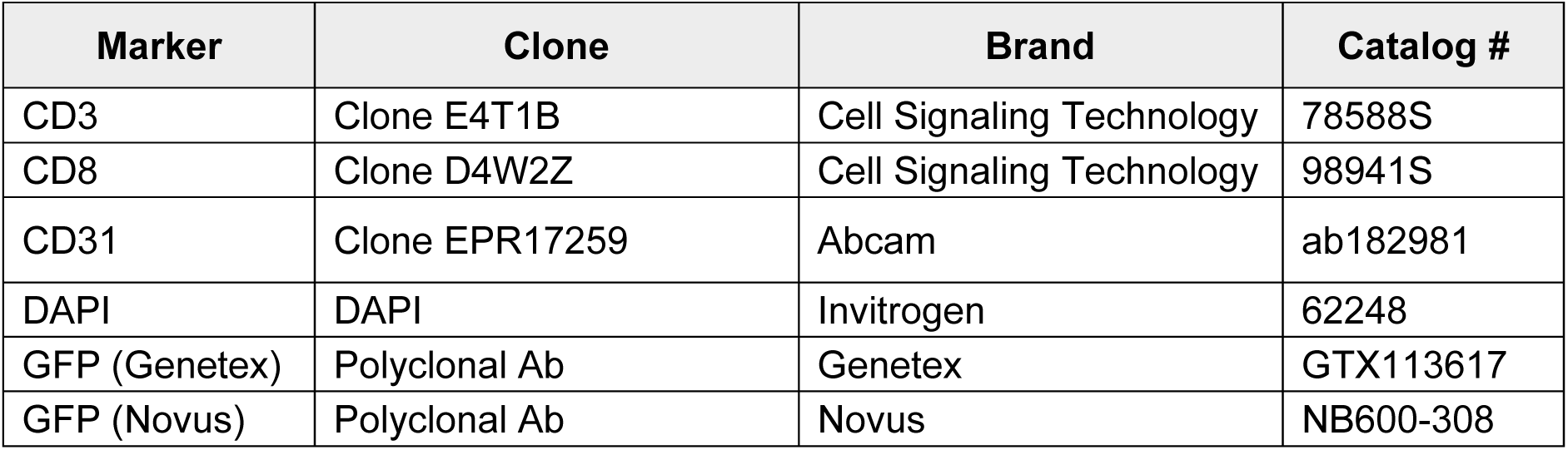
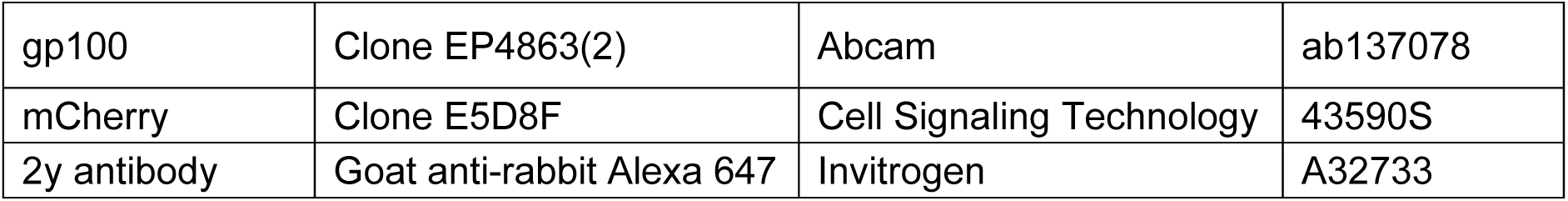

Briefly, the slides were deparaffinized, heat treated in antigen retrieval buffer, and serially incubated with primary antibodies followed by secondary antibodies conjugated to Alexa-647 (ThermoFisher). The slides were imaged using a 20x objective with 0.3 micron resolution and eluted between each staining round with Elution Buffer (Bio-techne). Consecutive images were aligned and autofluorescence and secondary-only staining background were subtracted using Horizon software. Imaris software (Oxford Instruments) was used to identify and create objects for each individual transferred T cell and measure their distance from the closest blood vessel in the tumor mass.

### Tumor survival studies

As described in tumor inoculation above, B78ChOva-hCD19 tumor cells were subcutaneously injected into one of the flanks of recipient CD45.1 mice or RAG mice. Five days later, activated CAR-T cells that overexpressed FMNL1 or control CAR-T cells were intravenously injected into separate cohorts of tumor bearing mice at 3.5 x 10^6^ cells/mouse.

Before CAR-T cell transfer, the recipient mice were randomized based on tumor size, with the control and FMNL1-overexpressing CAR-T cell groups having similar average initial tumor sizes. Recipient mice from both groups were co-housed. Tumor measurements were performed by personnel blinded to the type of CAR-T cells transferred. Tumor measurements were recorded daily with a manual caliper and tumor area was calculated in millimeters squared (length x width). Mice were monitored until IACUC-approved endpoint when tumor size was over 200 mm^2^, tumors had excessive ulceration, or mice lost 15% of their original body weight.

All mice were included in the analysis. After at least 55 days, mice in which tumors regressed and effectively disappeared were euthanized at the end of the experiment.

### Statistical analysis

Comparison of the means between samples was done using Graphpad Prism. *p-*values are denoted in the figure legends and in the figure images. All t-tests were two tailed and all analysis assumed both populations were normally distributed and parametric tests were used unless specified. Error bars are SEM unless otherwise indicated. The individual statistical tests used to analyze each experiment, and the experimental repeats, and sample sizes are reported in the figure legends.

## Results

### Establishment and characterization of FMNL1-overexpressing T cells

To investigate whether FMNL1 overexpression enhances T cell accumulation at tumor sites, we initially established a bioengineering platform to overexpress FMNL1 in T cells. We used ovalbumin-specific OT-I transgenic T cells and ovalbumin as a model antigen. OT-I T cells were activated and then transduced with retroviral constructs either co-expressing FMNL1 and GFP or a fluorescent protein only (such as Cerulean) as a control. The fluorescent proteins allowed us to sort T cells that integrated the retroviral construct and to track distinct cell populations in our experiments. In these experiments, FMNL1 protein expression was ∼1.8 fold higher in FMNL1-transduced T cells compared to control T cells, as determined by western blotting (**Fig. 1A**).

**Figure 1.**
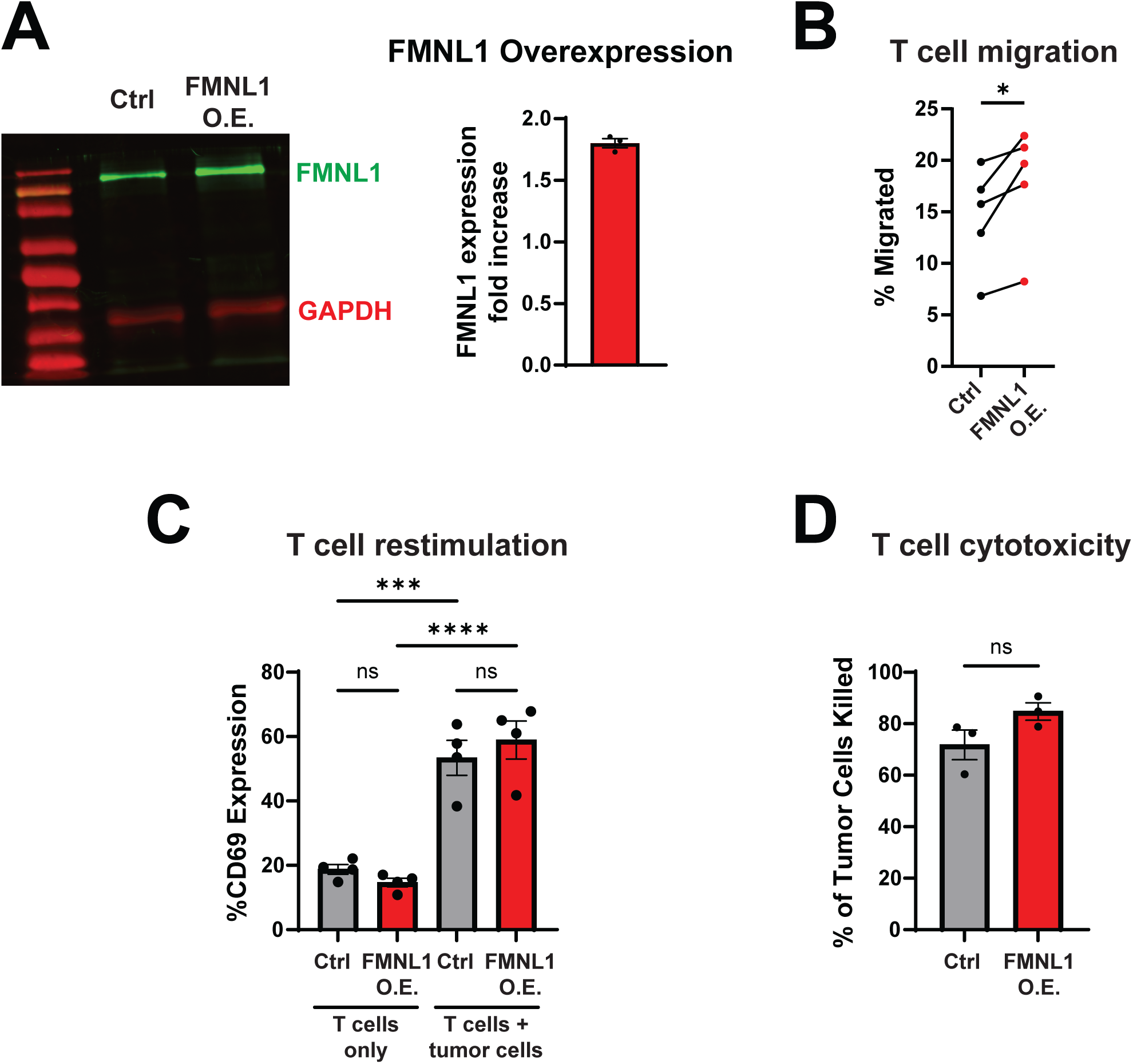
FMNL1-overexpressing T cells have increased migration through confining pores and normal T cell restimulation and killing capacity. **A)** T cells transduced with FMNL1 constructs have increased expression of FMNL1 protein compared to control transduced T cells. Left, representative western blot. Right, FMNL1 overexpression (O.E.) was confirmed by densitometry analysis of Western Blots. **B)** Overexpression of FMNL1 increases T cell migration through restrictive pores. Control and FMNL1-overexpressing T cells were allowed to migrate across 3μm pore transwell chambers with CXCL12 in the bottom wells for 1 hour and then quantified. **C-D)** FMNL1-overexpressing or control OT-I T cells were co-cultured with or without ovalbumin-expressing B78 melanoma cells for 24 hours and then assessed for CD69 expression (C) and tumor killing (D). In D the percent of tumor cells killed is calculated by comparing to a tumor cell only culture. Statistics were calculated using ordinary one-way ANOVA (C) or paired t-test (B, D), ns= not significant, *= p<0.05, ***= p<0.001, ****= p<0.0001. Data are the mean (±SEM) of ≥3 independent experiments.

We previously showed that FMNL1 *deficiency* does not alter *ex vivo* T cell activation, proliferation, and survival [32]. Here we sought to examine whether the *overexpression* of FMNL1 alters the phenotype of *ex vivo* activated T cells. First, we analyzed the expression of adhesion molecules (CD11a, CD49d) and chemokine receptors (CCR7, CXCR4, CXCR6) relevant for migration, as well as activation markers (CD44, CD62L, PD-1) We found no significant differences in the expression of most of these surface markers between FMNL1-overexpressing T cells and control T cells 5 days post-activation (Supplementary Fig. 1A-B). However, we detected a small but significant increase in CXCR6 expression in the FMNL1-overexpressing T cells (Supplementary Fig. 1B). CXCR6 plays a key role in T cell retention in solid tumors [8, 39, 40]. Additionally, T cells that receive strong activation have been suggested to upregulate CXCR6 in melanoma studies [8].

Our previous work demonstrated that FMNL1 plays a critical role in T cell migration through confining environments by facilitating nucleus deformation [29, 32]. We also found a linear correlation between FMNL1 expression levels and T cell transwell migration *in vitro* [32]. Given the key role of FMNL1 in T cell motility, we tested whether increasing FMNL1 expression could further enhance T cell migration through restrictive physical barriers. Using a reductionist transwell approach with 3µm pores to model restrictive barriers within solid tumors, we found that FMNL1-overexpressing T cells migrated significantly more than control T cells (**Fig. 1B**).

These findings support that FMNL1-mediated cytoskeletal remodeling enhances the capacity of T cells to traverse confined spaces, providing a potential mechanistic basis for improved tumor infiltration.

We next examined if FMNL1 overexpression would affect the ability of T cells to be restimulated by and kill target tumor cells using an *in vitro* co-culture system with OT-I T cells and ovalbumin-expressing melanoma cells. We found that FMNL1-overexpressing T cells had similar restimulation after 24 hours of co-culture with melanoma cells, as determined by CD69 expression (**Fig. 1C**). Furthermore, FMNL1-overexpressing OT-I T cells killed melanoma target cells expressing their cognate antigen as effectively as control OT-I T cells (**Fig. 1D**). These data support that FMNL1-overexpressing T cells have improved migration through confining pores while maintaining similar surface molecule profiles as control T cells. Furthermore, the overexpression of FMNL1 does not inhibit cell-cell interactions, restimulation, and cytotoxicity.

### FMNL1 overexpression promotes TIL accumulation at melanoma tumor sites

FMNL1 has a key role in T cell migration through confined collagen matrices [29] and we found that FMNL1 overexpression increases migration through restrictive pores *in vitro*. Thus, we next evaluated the impact of FMNL1 overexpression on T cell tumor infiltration *in vivo*. We subcutaneously implanted ovalbumin-expressing melanoma cells into the flanks of CD45.1-congenically marked recipient mice. Five days later, a timepoint when tumors are consistently palpable, control and FMNL1-overexpressing (CD45.2) OT-I T cells were co-transferred intravenously at a 1:1 ratio into recipient mice (**Fig. 2A**). Three and six days post T cell transfer, we quantified the number of transferred OT-I T cells at the tumor site by flow cytometry (**Fig. 2B**). We found significantly increased numbers of FMNL1-overexpressing OT-I T cells compared to control OT-I T cells at the tumor site at both timepoints post T cell transfer (**Fig. 2C**). While the total number of T cells in each individual tumor was variable, likely due to the varying tumor size and microenvironment, the ratio of FMNL1-overexpressing OT-I T cells compared to control OT-I T cells was consistently higher. Three days and six days post T cell transfer, there were respectively ∼1.6-fold and ∼1.7-fold more FMNL1-overexpressing T cells at the tumor site compared to control T cells (**Fig. 2C-D**).

**Figure 2.**
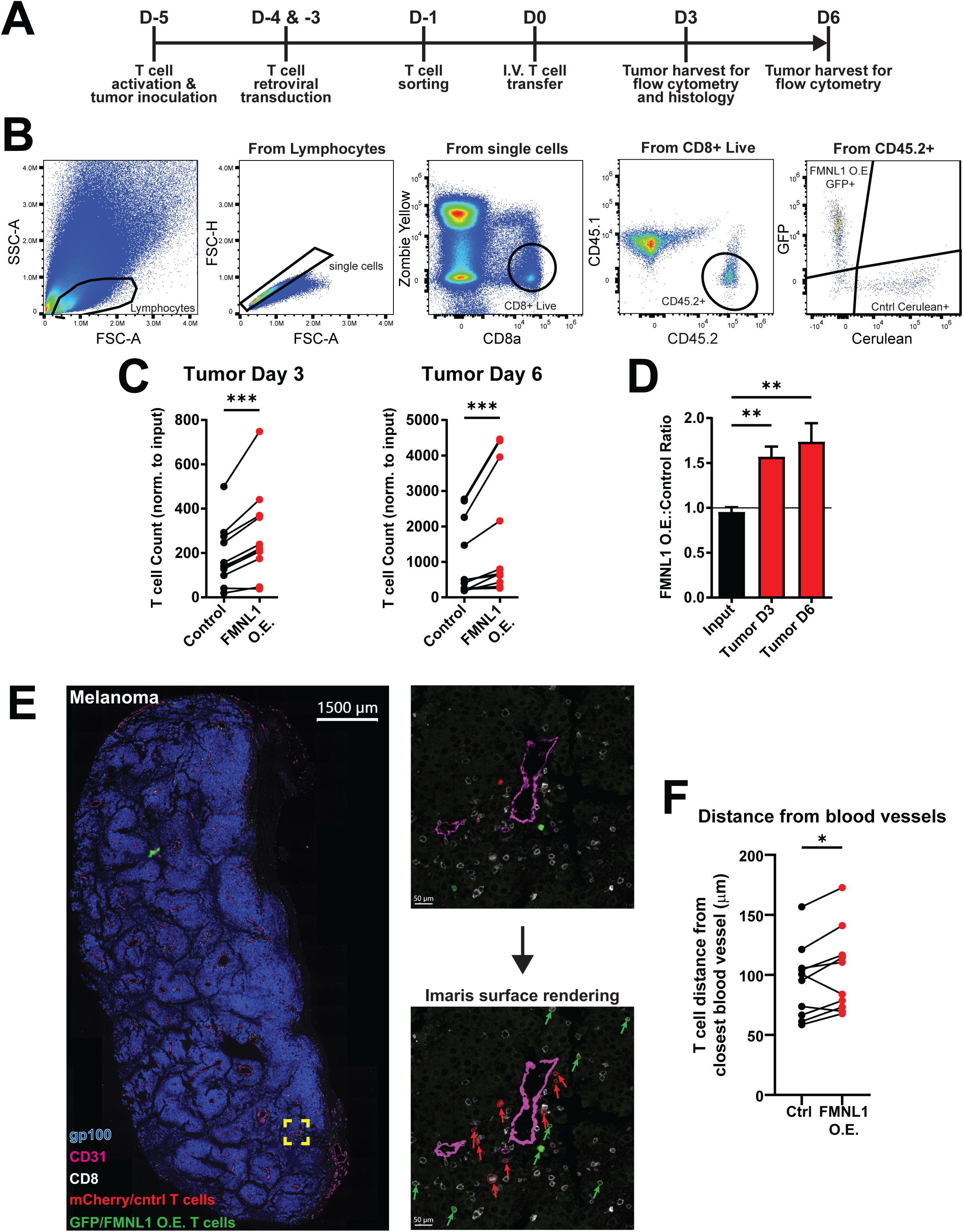
FMNL1 overexpression enhances TIL accumulation and infiltration at melanoma tumor sites. **A)** Schematic workflow for tumor inoculation, T cell activation and transduction, and tumor harvests. OT-I T cells were activated and transduced with FMNL1 or control vectors. Cells were sorted and a 1:1 mix of control and FMNL1-overexpressing (O.E.) T cells were injected into melanoma tumor bearing CD45.1 recipient mice. Tumors were collected 3-6 days post T cell transfer for evaluation by flow cytometry or histology. **B)** Representative flow cytometric analysis of TIL accumulation in tumors. Lymphocytes were gated by forward and side scatter and by a single cell gate (singlet). Then, transferred live T cells were gated using Live/Dead-, CD8+, CD45.1-, CD45.2+ gates. FMNL1-overexpressing cells were identified by GFP, and control cells were identified by Cerulean or mCherry. **C)** FMNL1 overexpression enhances TIL accumulation at the tumor site. Tumors were harvested 3 (left) or 6 days (right) post T cell transfer for quantification of transferred T cells by flow cytometry. The number of control and FMNL1-overexpressing T cells in each tumor is shown. Values were normalized to the injection ratio. **D)** Ratio of co-transferred FMNL1-overexpressing T cells to control T cells at the melanoma site on day 3 and day 6 post T cell transfer, compared to the measured injected ratio. A ratio above 1 indicates higher numbers of FMNL1-overexpressing T cells compared to control T cells. **E)** Tumors were collected 3 days post T cell transfer and processed for immunofluorescence staining. Left, representative image of stained whole tumor section showing tumor cells (gp100, blue), vascular endothelium (CD31, purple) CD8 T cells (CD8, white), FMNL1 O.E. OT-I T cells (GFP, green), control OT-I T cells (mCherry, red). Top right, magnified inset of the region indicated by the yellow square in the left image (gp100 staining is not shown for clarity). Bottom right, representative surface rendering of CD31+ blood vessels (purple), control OT-I T cells (CD8+/mCherry+, outlined in red) and FMNL1-overexpressing OT-I T cells (CD8+/GFP+, outlined in green) using Imaris software. **F)** FMNL1-overexpressing T cells have increased tumor infiltration. Quantification using Imaris software of the mean distance from the nearest blood vessel of control and FMNL1-overexpressing TILs. Statistics were calculated by paired ratio t test (in C, F) or ordinary one-way ANOVA (in D), *= p<0.05, **= p<0.01, ***= p<0.001. Data are from 3 independent experiments with ≥2 recipient mice/timepoint (in C, D) or 2 independent experiments each including at least 2 tumors/experiment and 2 sections/tumor (in F).

To determine whether FMNL1 enhanced TIL infiltration into the tumor parenchyma, we quantified the intratumoral spatial distribution of adoptively transferred OT-I T cells relative to tumor-associated blood vessels by immunofluorescence imaging. Control and FMNL1-overexpressing OT-I cells were co-transferred into ovalbumin-expressing melanoma bearing mice, and tumors were harvested 3 days later and analyzed by multi-spectral immunofluorescence imaging (**Fig. 2E**). FMNL1-overexpressing TILs were located significantly farther from tumor-associated blood vessels compared to control TILs (**Fig. 2F**), supporting increased migration away from the vasculature and deeper penetration into the tumor parenchyma compared to control TILs. These findings suggest that FMNL1-mediated cytoskeletal remodeling promotes T cell migration past the perivascular space and improves infiltration into tumor tissue.

We next assessed whether FMNL1 overexpression could inadvertently increase the accumulation of self-reactive T cells in healthy tissues expressing cognate antigen. To address this question, we used the pmel-1 transgenic TCR model, which recognizes the melanocyte antigen gp100/pmel-17 expressed by both melanoma cells and normal melanocytes. Consistent with our findings in the OT-I model, FMNL1 overexpression significantly increased pmel-1 T cell accumulation within melanoma tumors, resulting in ∼1.3-fold and ∼1.7-fold greater tumor accumulation at 2 and 6 days after adoptive transfer, respectively (**Fig. 3A-C**). Importantly, neither FMNL1-overexpressing nor control pmel-1 T cells showed substantial accumulation in normal skin, either adjacent to or distant from the tumor site (**Fig. 3A, C**). Furthermore, recipient mice did not develop vitiligo, a hallmark of melanocyte-directed off-tumor activity (not shown).

**Figure 3.**
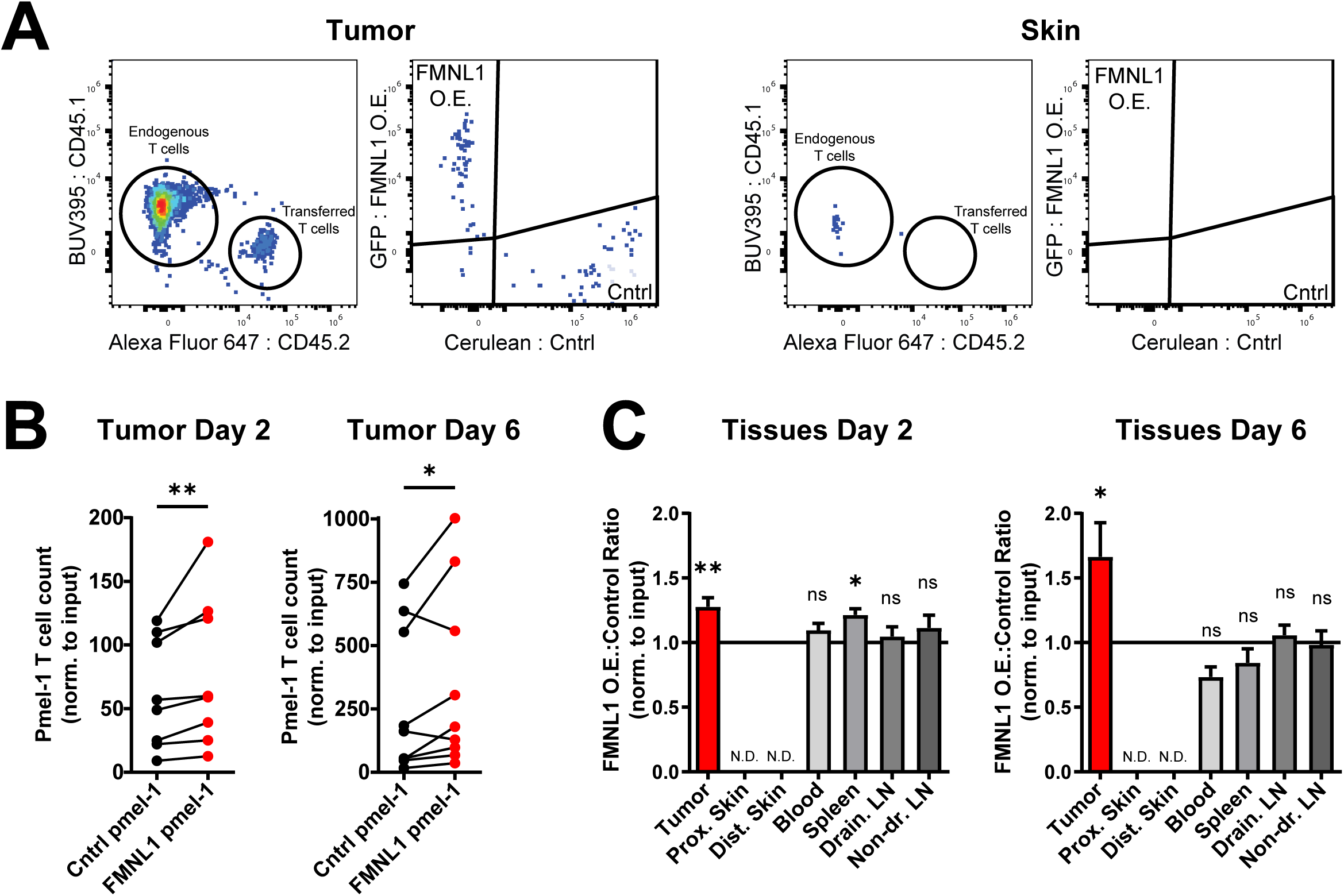
FMNL1 overexpression enhances pmel-1 TCR TIL accumulation at melanoma tumor sites but not in normal skin. FMNL1-overexpressing and control pmel-1 T cells were sorted and co-transferred at a 1:1 ratio into B16 melanoma bearing recipient mice. **A)** Representative flow cytometric analysis of T cell accumulation in tumors and skin. Lymphocytes were gated by forward and side scatter and by a single cell gate (singlet). Then, transferred live T cells were gated using Live/Dead-, CD8+, CD45.1-, CD45.2+ gates. FMNL1-overexpressing cells were identified by GFP and control cells were identified by Cerulean. **B)** Number of control and FMNL1-overexpressing pmel-1 T cells at the melanoma site on day 2 (left) or day 6 (right) post T cell transfer. Values were normalized to the injection ratio. **C)** Ratio of co-transferred FMNL1-overexpressing to control pmel-1 T cells at the melanoma site and at other non-tumor tissues 2 days (left) or 6 days (right) post T cell transfer. A ratio above 1.0 (black line) indicates an increase in FMNL1-overexpressing cells. Ratios in the skin tissues were not determined given that transferred T cells were generally below level of detection. Data are from 3 independent experiments each with ≥1-2 recipient mice/timepoint. Statistics were calculated by ratio paired t test (B) or using a one-sample t-test against a theoretical FMNL1 OE:Control ratio of 1.0 (C). ns= not significant, *= p<0.05, **= p<0.01.

These findings support that FMNL1 overexpression enhances tumor-reactive T cell accumulation and persistence at the tumor site without substantial infiltration in antigen-expressing healthy tissues or increased off-tumor toxicity.

### FMNL1-overexpressing CAR-T cells have increased accumulation at the tumor site

Having determined that FMNL1 overexpression can enhance TIL accumulation at melanoma sites, we wanted to confirm and expand our findings in a CAR-T cell system. For these experiments, we used a well-established CAR construct that targets human CD19 (hCD19) and expressed hCD19 in melanoma cells as a model tumor antigen [38, 41]. First, to engineer CAR-T cells, we activated WT CD8 T cells with anti-CD3/anti-CD28 coated beads and then transduced them with either anti-hCD19 CAR and FMNL1 or anti-hCD19 CAR and control retroviral vectors. For these experiments, we sorted for, and utilized, T cells expressing high levels of the GFP fluorescent marker co-expressed on the FMNL1 vector, hypothesizing that this would lead to greater FMNL1 overexpression. Control CAR-T cells were separately sorted based on the Cerulean fluorescent marker on the control vector. By western blotting, we observed a high level of FMNL1 overexpression in CAR-T cells, which on average was ∼3-fold higher than control transduced CAR-T cells (Supplementary Fig. 2A). We then examined whether the overexpression of FMNL1 would alter the CAR-T cell phenotype from effector to memory or alter chemokine receptor expression when activated *ex vivo* [41, 42]. We found no significant difference in CD122 (IL-2Rβ), CD127 (Il-7Rα) and chemokine receptors CXCR4 and CXCR6, suggesting that FMNL1-overexpressing CAR-T cells have similar expression profiles as control CAR-T cells (Supplementary Fig. 2B-C).

Next, to evaluate CAR-T cell accumulation at tumor sites, we co-transferred sorted FMNL1-overexpressing CAR-T cells and control CAR-T cells at a 1:1 ratio into congenically-marked CD45.1 mice bearing hCD19-expressing melanoma tumors. Three or six days later, we quantified the number of transferred FMNL1-overexpressing and control CAR-T cells in tumors, as well as blood, spleen, draining LN, and non-draining LN of recipient mice following a similar gating scheme as in Figure 2 with the addition of CD90.1/Thy1.1 (expressed on the CAR vector) to identify the CAR^+^ T cells (**Fig. 4A)**. Our data showed a significantly increased number of FMNL1-overexpressing CAR-T cells compared to control CAR-T cells at the melanoma tumor site (**Fig. 4B**). We observed a ∼1.8-fold increase in FMNL1-overexpressing CAR-T cells in the tumor 3 days post-transfer and a ∼3.8-fold increase 6 days post-transfer compared to control CAR-T cells (**Fig. 4B-C**). Analysis of secondary lymphoid organs (SLOs) 3 days post transfer suggested slightly increased numbers of CAR-T cells that overexpress FMNL1 in SLOs.

**Figure 4.**
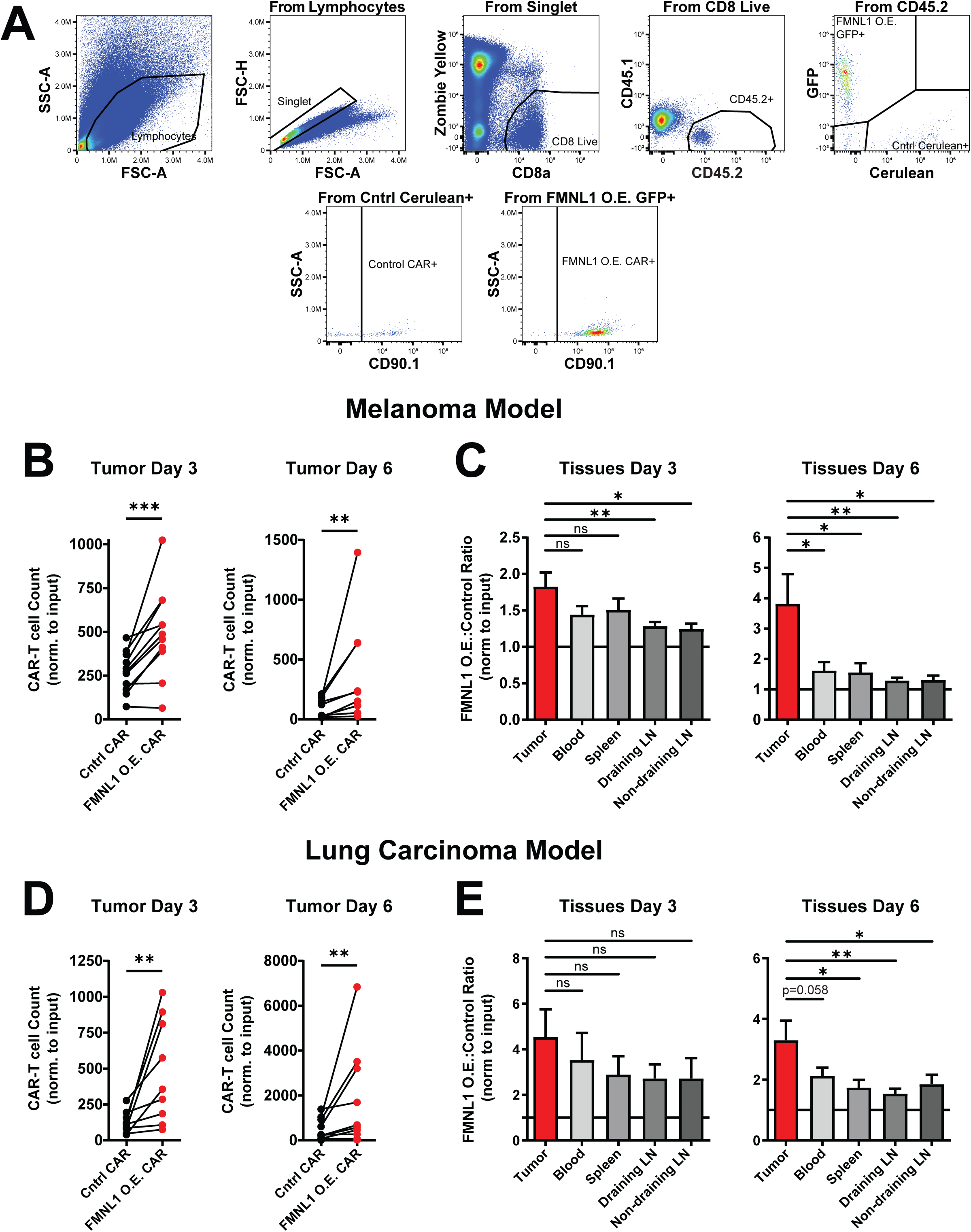
FMNL1 overexpression enhances accumulation and persistence of CAR-T cells at solid tumor sites. Transduced FMNL1-overexpressing anti-hCD19 CAR-T cells and control anti-hCD19 CAR-T cells were co-injected at a 1:1 ratio into hCD19-expressing tumor bearing CD45.1 recipient mice. Tumors were harvested 3 or 6 days post CAR-T cell transfer for quantification by flow cytometry. **A)** Representative flow cytometric analysis of FMNL1-overexpressing (O.E.) and control CAR-T cell accumulation in tumors. Transferred live T cells were gated as in figure 2, with FMNL1 overexpressing cells identified by GFP and control cells identified by Cerulean. CAR-T cells were further identified by CD90.1. **B)** FMNL1 overexpression enhances CAR-T cell accumulation at melanoma tumor sites. The number of FMNL1-overexpressing CAR-T cells and control CAR-T cells at the B78 melanoma tumor site is shown. **C)** Ratio of co-transferred FMNL1-overexpressing CAR-T cells to control CAR-T cells at the melanoma site and other non-tumor tissues on day 3 (left) or day 6 (right) post CAR-T transfer. A ratio above 1 (black line) indicates higher numbers of CAR-T cells that overexpress FMNL1 compared to control CAR-T cells. Values were normalized to the injection ratio. **D)** FMNL1-overexpressing and control CAR-T cells were co-transferred into hCD19-expressing LL/2 lung carcinoma bearing CD45.1 recipient mice. The number of FMNL1-overexpressing and control CAR-T cells at the ectopic lung carcinoma site is shown. **E)** Ratio of co-transferred FMNL1-overexpressing CAR-T cells to control CAR-T cells T cells at the lung carcinoma tumor site and other non-tumor tissues on day 3 (left) or day 6 (right) post CAR-T transfer. Data in E is depicted as described in C. Statistics were calculated by ratio paired t test (B, D) or by ordinary one-way ANOVA (C, E), ns= not significant, * = p<0.05, **= p<0.01, ***= p<0.001. Data are the mean (+SEM) of at least 3 independent experiments with 2 recipient mice/timepoint.

However, when comparing the ratio of FMNL1-overexpressing to control CAR-T cells in the tumor to the ratio in other tissues, by 6 days post CAR-T cell co-transfer, FMNL1-overexpressing CAR-T cells had a significantly and selectively higher ratio in the tumor than all the other tissues analyzed (**Fig. 4C**). This supports that over time FMNL1 overexpression leads to CAR-T cell accumulation and persistence specifically at the tumor site over SLOs and blood.

We then investigated if overexpression in CAR-T cells of mDia1, the other Formin highly expressed in T cells, would have a similar effect on CAR-T cell accumulation in melanoma tumors. Using a similar experimental setup as for the FMNL1 overexpression experiments, we found enhanced accumulation of mDia1-overexpressing CAR-T cells selectively at the melanoma tumor site (Supplementary Fig. 3). These data suggest that, similar to FMNL1, mDia1 overexpression in CAR-T cells can promote trafficking to melanoma sites and increases accumulation of CAR-T cells at the tumor site.

We then wanted to determine whether our findings on FMNL1 overexpression enhancing CAR-T cell accumulation in melanoma extended to other solid tumor models. Another “cold” tumor model with low T cell infiltration and high presence of suppressive myeloid cells is the Lewis lung carcinoma (LL/2) model [43, 44]. This LL/2 model recapitulates human lung adenocarcinoma genomically with similar oncogenes [45]. We expressed hCD19 in LL/2 cells and then ectopically injected LL/2 cells expressing hCD19 into the flanks of recipient mice and 5 days later co-transferred a 1:1 ratio of FMNL1-overexpressing CAR-T cells and control CAR-T cells. Similar to the melanoma model, we observed significantly more FMNL1-overexpressing CAR-T cells compared to control CAR-T cells at the tumor site on both day 3 and day 6 post CAR-T cell transfer (**Fig. 4D**). There were ∼4.5-fold and ∼3.3-fold increases in tumor accumulation of FMNL1 overexpressing CAR-T cells compared to control CAR-T cells on day 3 and day 6 respectively (**Fig. 4D-E**). Additionally, we found that by day 6, the ratio of FMNL1-overexpressing CAR-T cells to control CAR-T cells at the LL/2 lung carcinoma site was significantly higher than the ratio in the other SLOs analyzed (**Fig. 4E**). Given the similar effect in two different cancer models tested, our data supports that FMNL1 overexpression increases CAR-T cell accumulation and persistence at solid tumor sites.

### Adoptive transfer of FMNL1 overexpressing CAR-T cells limits tumor growth and improves survival of tumor bearing mice

Having seen higher CAR-T cell accumulation in tumors, we investigated if the enhanced accumulation of FMNL1-overexpressing CAR-T cells at the tumor site would limit tumor growth and improve survival of tumor-bearing recipient mice. For these experiments, we transferred FMNL1-overexpressing or control CAR-T cells into hCD19-expressing melanoma bearing RAG knock-out recipient mice to avoid potential CAR-T cell rejection and to mimic the use of lymphodepletion prior to CAR-T transfer [46, 47]. Our data showed significantly prolonged survival of recipient mice that received FMNL1 overexpressing CAR-T cells (**Fig. 5A**).

**Figure 5.**
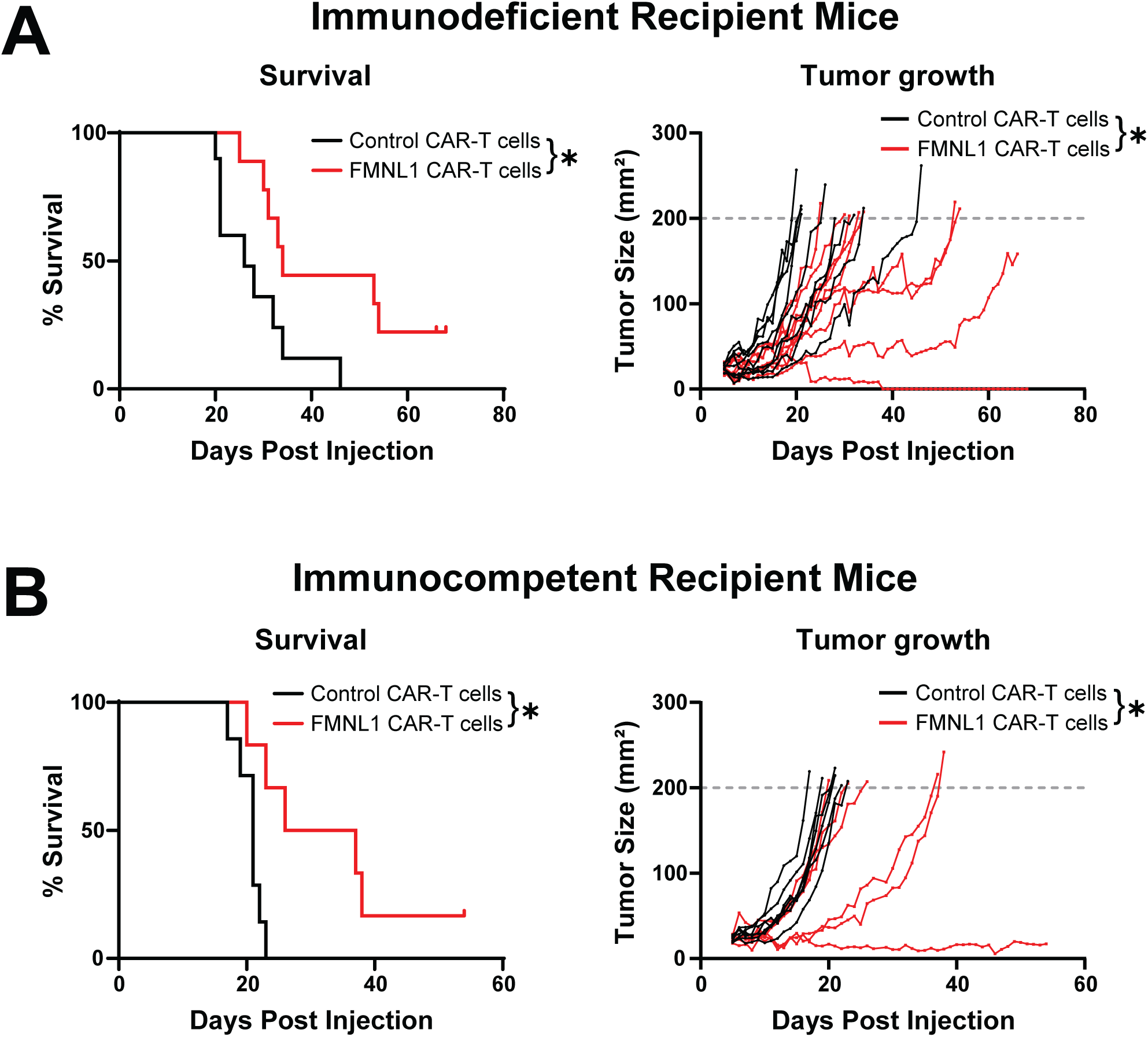
FMNL1-overexpressing CAR-T cells improve overall survival of tumor bearing mice. FMNL1-overexpressing or control anti-hCD19 CAR-T cells were transferred into hCD19^+^ B78 melanoma bearing RAG knock-out mice (in **A**) or immunocompetent mice (in **B**). Tumor growth was monitored daily and mice were euthanized if tumor size exceeded a 200 mm^2^ cutoff. Survival curves (left) and individual tumor sizes (right) are shown. Statistics were calculated by Log-rank test for the survival curves and mixed-effects model for tumor growth comparisons. * = p<0.05. Data in A are pooled from 3 independent experiments with 2-4 mice/group/experiment. Data in B are pooled from 2 independent experiments, each with cohorts of 3-4 mice/group/experiment.

Furthermore, compared to tumor bearing mice adoptively transferred with control CAR-T cells, mice receiving FMNL1-overexpressing CAR-T cells showed delayed tumor growth and in one case complete tumor regression and eradication (**Fig. 5A**). Next, we explored if FMNL1-overexpressing CAR-T cells could also enhance survival in immunocompetent mice. Similar to the survival studies in RAG knock-out recipient mice, immunocompetent tumor-bearing mice had delayed tumor growth and survived significantly longer when treated with FMNL1-overexpressing CAR-T cells compared to control CAR-T cells (**Fig. 5B)**. Overall, these data show that, compared to control CAR-T cells, CAR-T cells that overexpress FMNL1 significantly limit tumor growth and improve overall survival of melanoma-bearing mice. Importantly, we saw this effect in both immunodepleted and immunocompetent host settings, suggesting that the effect of FMNL1 overexpression is not dependent on an immune depleted environment in the recipient host.

## Discussion

Overall, our findings demonstrate that overexpression of FMNL1 in TILs and CAR-T cells increases intratumoral T cell accumulation, limits/delays tumor growth, and prolongs survival of tumor-bearing mice. Both FMNL1 and mDia1 overexpression enhanced CAR-T cell accumulation at the solid tumor site. This finding is consistent with the established roles of FMNL1 and mDia1 in transendothelial migration and trafficking to inflamed tissues [32, 34].

Furthermore, we have recently shown that FMNL1 and mDia1 have distinct subcellular localization patterns and roles in T cell migration [29]. Specifically, we determined that FMNL1 is necessary to promote nuclear deformation and passage through dense, restrictive 3D collagen matrices, while mDia1 is important for 3D motility in general [29]. Here, we found that FMNL1 overexpression improved T cell migration through confining pores *in vitro* and increased the depth of TIL tumor infiltration *in vivo*. Thus, these findings support a model in which overexpression of FMNL1 promotes T cell migration across the abnormal tumor vasculature and through the dense and fibrotic ECM that surrounds many solid tumors. Together, our previous and current results suggest that improved migration through the restrictive tumor environment likely contributes, at least in part, to the enhanced accumulation of FMNL1-overexpressing T cells within tumors.

Interestingly, FMNL1-overexpressing T cells progressively accumulated more specifically at the tumor site compared to other non-tumor tissues over time. This finding suggests that increased accumulation may represent a combination of enhanced initial infiltration and of improved persistence due to increased proliferation, survival, or retention in response to antigen encounter in the TME. Additional investigation will be needed to fully determine the mechanisms that mediate the enhanced accumulation and the persistence of FMNL1-overexpressing T cells at tumor sites. Future studies combining spatial and phenotypic characterization of transferred T cells could help distinguish the relative contributions of these processes and provide mechanistic insight into how FMNL1-driven cytoskeletal remodeling promotes durable T cell accumulation within tumors and effective antitumor immunity.

Although FMNL1-overexpressing CAR-T cells significantly delayed tumor growth and extended survival of tumor bearing mice, tumors were not uniformly eliminated. This incomplete response suggests that FMNL1-overexpressing CAR-T cells may become partially exhausted or dysfunctional within the tumor microenvironment. Additionally, tumor escape through loss of target antigen expression may also contribute to eventual disease progression. These findings highlight the potential value of combining FMNL1 overexpression with next generation CAR designs, immune checkpoint inhibitors, and/or strategies to prevent tumor antigen loss to further improve the therapeutic efficacy of these individual approaches.

Overall, this work validates cytoskeletal engineering through FMNL1 overexpression as a novel approach to enhance T cell infiltration capabilities through abnormal tissue architecture in the TME, increase T cell persistence at the tumor site, and improve therapeutic activity in solid tumors. Importantly, this approach does not depend on a specific CAR or TCR construct, or MHC restriction. Notably, we observed enhanced accumulation of FMNL1-engineered T cells in solid tumors in both immunocompetent and immunodeficient settings, supporting broad applicability of this approach to a variety of therapeutic scenarios and treatments. Together, these findings identify FMNL1 as a cytoskeletal engineering target for overcoming infiltration barriers in the TME and support the integration of FMNL1 engineering with existing and emerging T cell-based immunotherapies for the treatment of solid tumors.

## Acknowledgements

We thank B. Basta for help with mouse genotyping and colony maintenance; S. Beard for technical assistance with cell sorter and flow cytometer maintenance; K. Kremer, R. Friedman, M. Verneris, and Y. Zhu for their feedback during the manuscript writing process. We also thank the University of Colorado OLAR personnel for animal husbandry and technical assistance.

## Funding

This work was supported in part by grants from: Gates Institute Grubstake award (JJ), NCI R21CA280145 (JJ), and NIAID R01AI167943 (JJ). This work was also supported in part by the University of Colorado Diabetes Research Center (DRC) grant P30DK116073 and the Human Immune Monitoring Shared Resource (HIMSR) at the University of Colorado School of Medicine was supported in part by NIH S10OD036353. The content of this work is solely the responsibility of the authors and does not necessarily represent the official views of the NIH.

## Authors’ contributions

JWC conceptualized the studies, performed experiments and data analysis, and wrote the paper. JO-C performed experiments and data analysis. AMW performed experiments and data analysis. HS performed experiments and data analysis. ALS performed experiments and data analysis. EN performed experiments. MEA performed experiments. KHT performed experiments. JJ conceptualized, supervised and acquired funding for the studies, and wrote the paper.

## Data Availability Statement

The data from this study are either included in the article or in supplementary information, are publicly available through ImmPort study SDY3641 (DOI:10.21430/M34PX11DIU), or will be made available upon reasonable request.

## Conflict of interest disclosure statement

The University of Colorado has a patent application related to this study on which JJ is named as an inventor.

---

**Supplementary Figure 1. *Ex vivo* activated FMNL1-overexpressing and control OT-I T cells express similar levels of surface molecules and chemokine receptors. A)** Representative overlay histograms of antibody staining for the indicated surface molecules quantified in B. **B)** Ratio of geometric mean fluorescent intensity (gMFI) of antibody staining of indicated surface molecules on FMNL1-overexpressing (O.E.) T cells compared to control T cells 5 days after activation. A ratio above or below 1.0 (solid line) indicates a respective increase or decrease in expression of indicated surface molecules on FMNL1-overexpressing T cells relative to control T cells. Data are from ≥3 independent experiments (±SEM). Statistics were calculated using a one-sample two-tailed t-test against a theoretical FMNL1 OE:Control gMFI ratio of 1.0. n.s.= not significant, * = p<0.05.

**Supplementary Figure 2. *Ex vivo a*ctivated FMNL1-overexpressing and control CAR-T cells express equivalent levels of cytokine and chemokine receptors. A)** Representative western blot showing FMNL1 overexpression (O.E.) in anti-hCD19 CAR-T cells (left). FMNL1-overexpression was analyzed by densitometry analysis of Western Blots (right). **B)** Representative overlay histograms of antibody staining for the indicated surface molecules quantified in C. **C)** Ratio of geometric mean fluorescent intensity (gMFI) of antibody staining of indicated surface molecules of FMNL1 overexpressing CAR-T cells compared to control CAR-T cells 5 days after activation. A ratio above or below 1.0 (solid line) indicates a respective increase or decrease in expression of indicated surface molecules on FMNL1-overexpressing CAR-T cells relative to control CAR-T cells. Data are the mean (±SEM) from ≥4 independent experiments. Statistics were calculated using a one-sample two-tailed t-test against a theoretical FMNL1 OE:Control gMFI ratio of 1.0. n.s.= not significant.

**Supplementary Figure 3. mDia1-overexpressing CAR-T cells have increased numbers at melanoma tumor sites.** mDia1-overexpressing (O.E.) anti-hCD19 CAR-T cells or control anti-hCD19 CAR-T cells were co-transferred at a 1:1 ratio into mice bearing hCD19-expressing B78 melanoma tumors. The tumors were then harvested 6 days post CAR-T cell transfer for quantification of CAR-T cells by flow cytometry. **A)** Quantification of control and mDia1-overexpressing CAR-T cells at the tumor site 6 days post CAR-T cell transfer. The number of mDia1-overexpressing CAR-T cells and control CAR-T cells at the melanoma tumor site is shown. **B)** Ratio of co-transferred mDia1-overexpressing CAR-T cells to control CAR-T cells in the indicated tissues on day 6 post CAR-T cell transfer. Values were normalized to the injection ratio. A ratio above 1 (black line) indicates higher numbers of CAR-T cells that overexpress mDia1 compared to control CAR-T. Statistics in A were calculated using ratio paired t test. Statistics in B were calculated by one-sample two-tailed t-test against a theoretical mDia1 O.E:control ratio of 1.0. ns=not significant, * = p<0.05, **= p<0.01. Data are the mean (+SEM) of 2 independent experiments.

## Notes

https://immport.org/shared/study/SDY3641/summary

### Competing Interest Statement

The University of Colorado has a patent application related to this study on which Jordan Jacobelli is named as an inventor.

