## Supplementary figures and images for "Cytoskeletal engineering through Formin-like 1 overexpression enhances T cell infiltration and antitumor potency in solid tumors"

### Supplemental Figure 1

*Chung et al. Supplementary Figure 1*

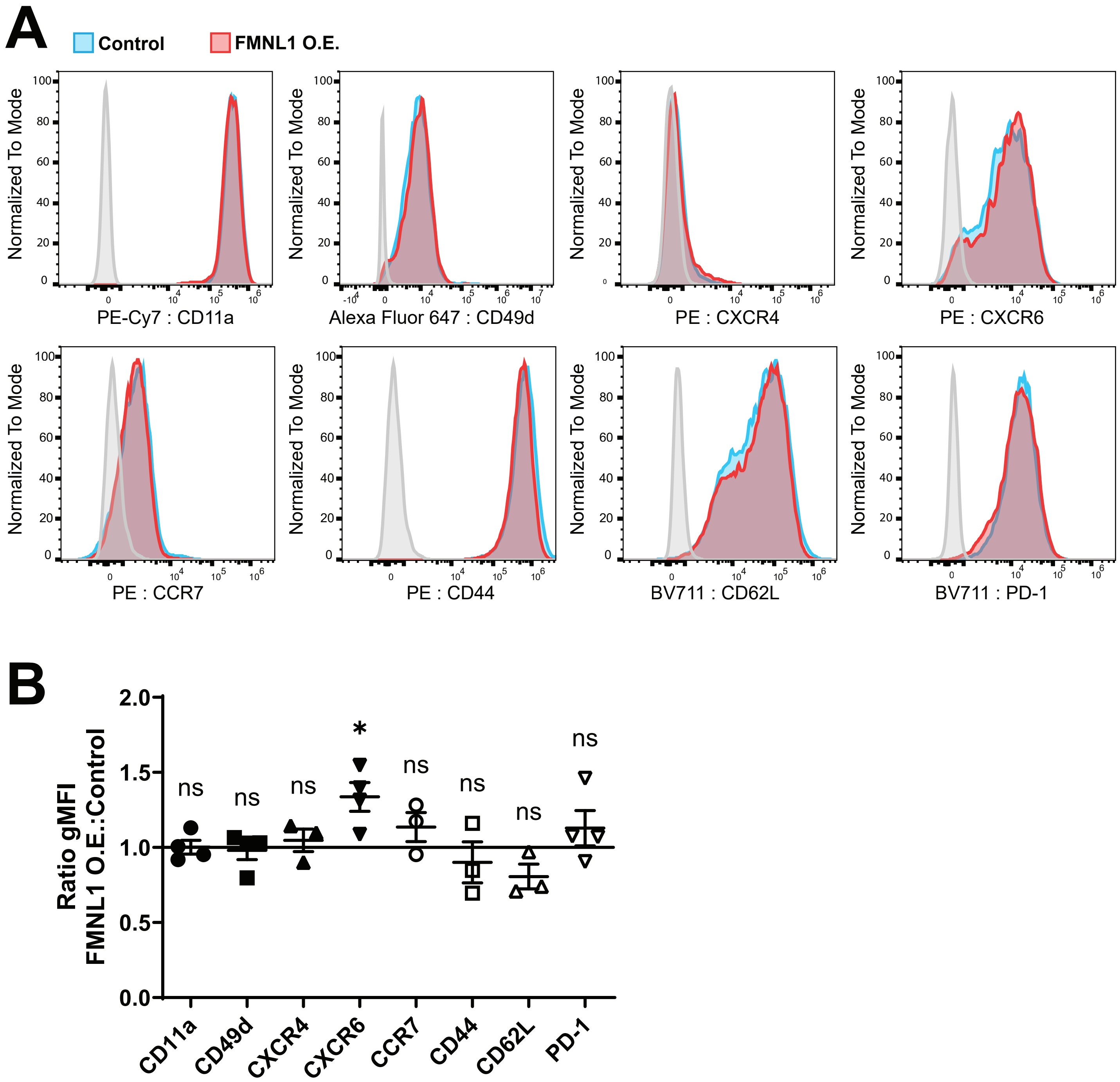

### Supplemental Figure 2

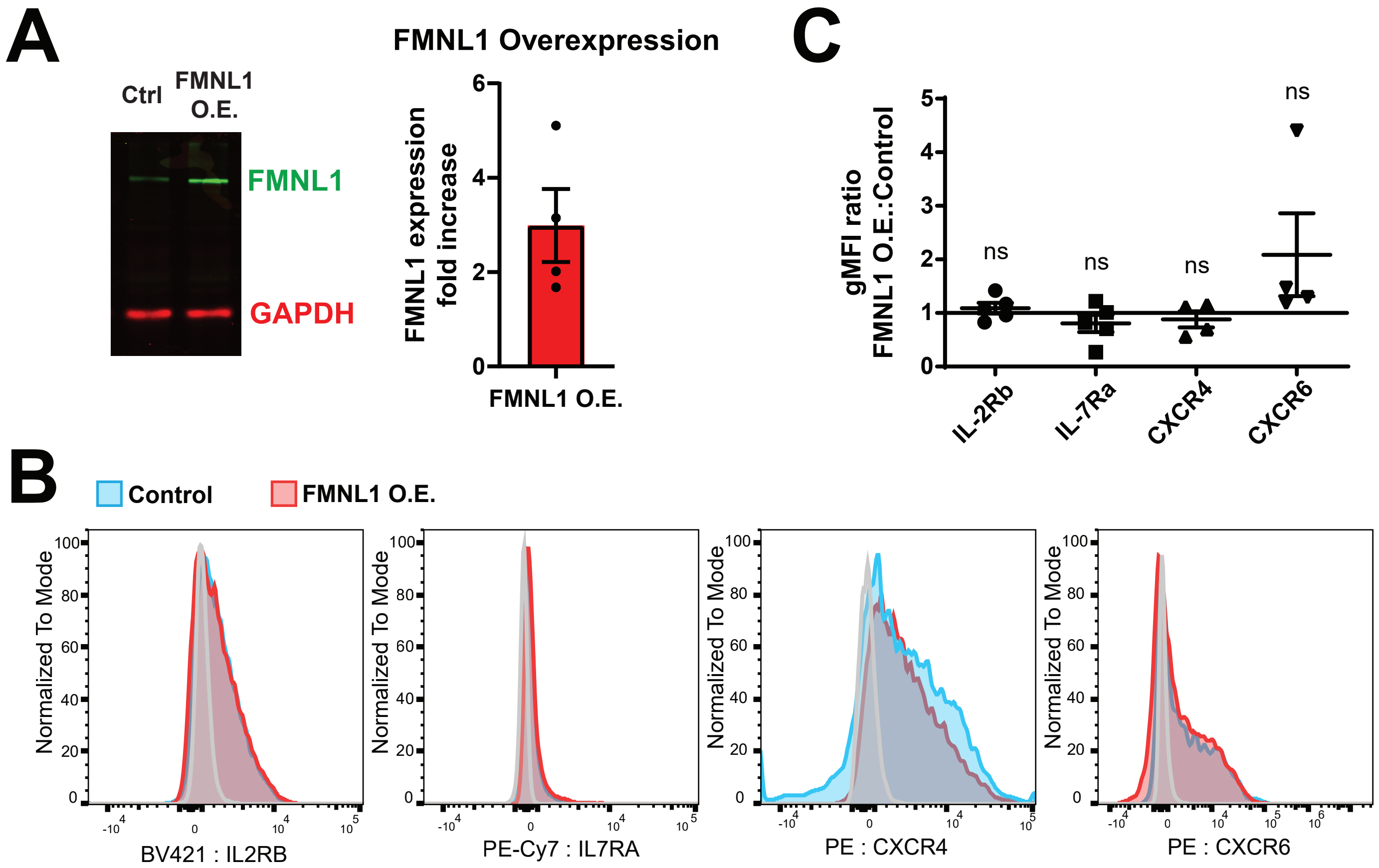

### Supplemental Figure 3

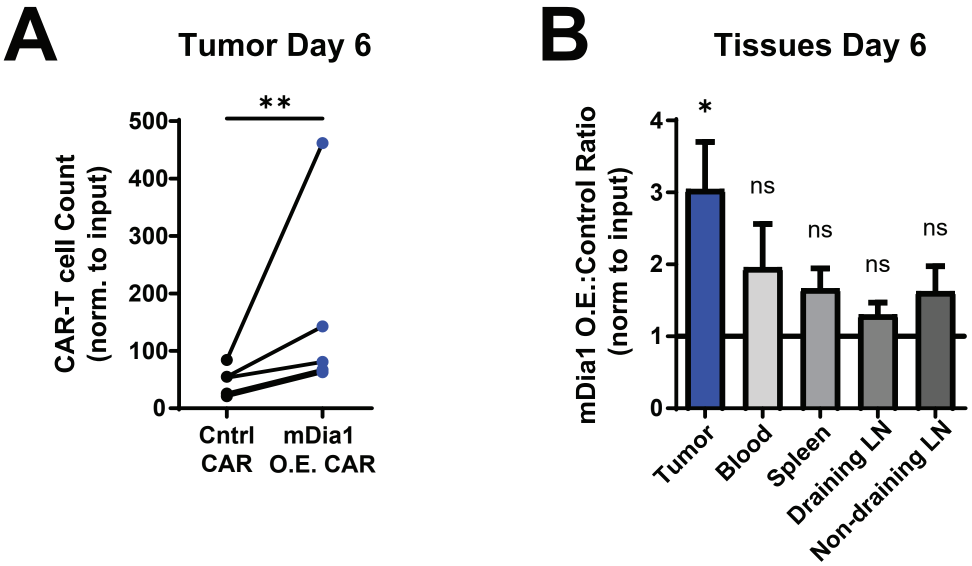
